# QutRNA2: Robust tRNA modification discovery from Nanopore direct tRNA sequencing

**DOI:** 10.1101/2025.10.20.683443

**Authors:** Michael Piechotta, Wei Guo, Isabel S. Naarmann-de Vries, Felix Kallenborn, Francesca Tuorto, Bertil Schmidt, Christoph Dieterich

## Abstract

Transfer RNAs (tRNAs) are essential for protein synthesis and are extensively modified to ensure their structure and function. Direct RNA sequencing with Oxford Nanopore Technologies enables positional modification analysis but is challenged by tRNAs’ short length, redundancy, and dense modifications. We present QutRNA2, a scalable workflow featuring GPU-accelerated local alignment, statistical filtering, pairwise error profile comparison, and customizable visualization. Achieving up to 25-fold speed gains over CPU methods, QutRNA2 identifies enzyme-dependent modifications in nuclear- and mitochondrial-encoded tRNAs, demonstrated in human and mouse samples. This open-source solution provides a comprehensive, multiplexing-compatible framework for tRNA analysis, addressing a key gap in current tools.

## 1 Background

Transfer RNAs (tRNAs) play a central role in protein translation by decoding mRNA codons and providing amino acids to the ribosomes. Chemical modifications heavily influence tRNA folding, stability, and codon-anticodon pairing [1]. More than 170 modifications have been identified, and the number is increasing as the development of new detection methods continues [2]. Each tRNA is typically modified at 5-15 positions (average 8) [3]. Most modifications are evolutionarily conserved [4], with increasing complexity in terms of the number of modifications and the chemical structure of the modifications from simpler organisms like *E*.*coli* to more complex eukaryotes as humans [2].

Recently, tRNAs have been sequenced using Oxford Nanopore Technologies (ONT) direct RNA sequencing chemistry [5–7]. Here, modified residues alter current signal traces and lead to out-of-domain signals, which in turn result in basecalling errors that manifest as substitutions, insertions, or deletions of canonical RNA bases. The comparison of these error profiles has been used for the identification of RNA modifications for multiple RNA types in the past [8]. The unique properties of tRNAs, such as their short length, the presence of several identical copies in the genome, and the high frequency of modifications, pose a challenge to the mapping algorithm employed. Traditional approaches, such as seed-extend methods, often fail to achieve satisfactory accuracy.

We have recently presented an improved alignment protocol and modification detection workflow (coined QutRNA) that successfully identified tRNA queuosine modifications in tRNAs of *Schizosaccharomyces pombe* and *Escherichia coli* [9].

Here, we present an improved and extended workflow (QutRNA2: https://github.com/dieterich-lab/QutRNA2), which masters a much higher transcriptome complexity, sequencing throughput and the new RNA004 chemistry. Briefly, QutRNA2 involves four principle components (see Figure 1A): (1) an optimal local GPU-accelerated ultra-fast read alignment step, Figure 1B, (2) rigorous statistical filtering of spurious alignments, (3) post selection of aligned reads by formal requirements (e.g. adapter overlap) (4) a new score normalization approach by subsampling. The reported scores are robust to technical fluctuations and serve to highlight candidate modification positions in the Sprinzl coordinate space [10] (see Figure 1G).

**Fig. 1:**
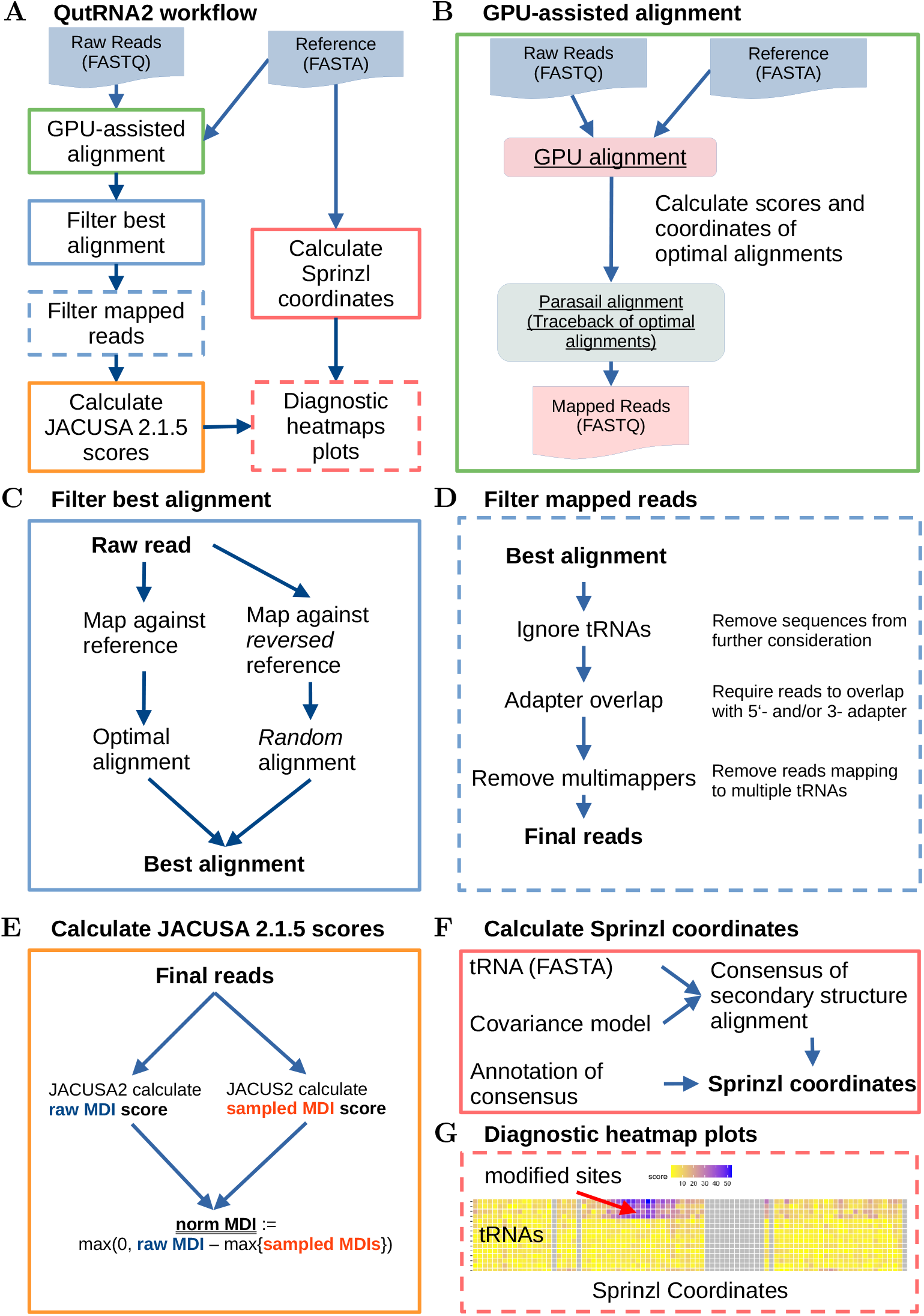
(A) Summary of the QutRNA2 workflow to generate diagnostic heatmap plots from raw reads and reference. (B) GPU-accelerated alignment of raw reads. First, optimal alignments are identified with the ultra-fast GPU alignment. Next, the coordinates are expanded with parasail to generate an alignment traceback, alignment score, and SAM output. (C) Best alignments are filtered against a random alignment by repeating the alignment for all reads against the reversed reference sequence. (D) Filtering mapped reads. A set of possible filters with respective descriptions. The filters are optional, and the order of their application is arbitrary. (E) Calculate JACUSA2 scores that are indicative of RNA modifications. JACUSA2 scores are normalized by subsampling reads (see Supplementary Figure S2 for details). (F) Calculate Sprinzl coordinates. Reference sequence and matching covariance model are used to calculate secondary structure alignments. Finally, the resulting consensus alignments are labeled, producing the Sprinzl coordinates. (G) Diagnostic heatmap plots. Subsampled JACUSA2 scores are plotted for tRNAs (rows) and Sprinzl coordinates (columns). High scores indicate putative modification sites.

JACUSA2 [11] remains at the heart of QutRNA2 for modification detection by pairwise comparisons. JACUSA2 detects RNA modifications by calculating and comparing error profiles derived from mismatches, insertions, and deletions in paired samples from control, knock down, knock out or IVT conditions for a wide range of modification types [12].

We demonstrate the performance of QutRNA2 in HCT116 cells harboring knock outs of known tRNA-modifying enzymes such as QTRT1 and NSUN2 in nuclear and mitochondrial tRNAs. Finally, we extend the analysis to experiments in *Mus musculus* where we study the impact of the absence of Qtrt1 protein on tRNA modifications in murine embryonic stem cells (mESCs) and hippocampus (mHC).

## 2 Results

### 2.1 QutRNA2 features and impact

We have expanded, added, and enhanced several features of the QutRNA workflow to meet new challenges in data volume and complexity as outlined below (see Background and Figure 1).

#### 2.1.1 Rapid GPU-accelerated optimal local alignment

In our previous work [9], we have shown that optimal local read alignments are superior to heuristic approaches, such as the classical seed-and-extend read mapping concept. The latter often mis-assigns reads to false database records. However, data processing time becomes prohibitively high if millions of reads need to be analyzed. Figure 2A compares the running time for mapping 1 million reads against a human reference tRNome using our previous approach and our newly implemented GPU-accelerated read mapper (see Methods 5.5). We achieve a median 25-fold speedup. Our subsequently presented and discussed tRNA RNA004 libraries comprise ≈ 20 million reads. Each read must be mapped twice to create random alignments to deduce an appropriate alignment score threshold. Therefore, ≈ 40 million reads must be mapped to the database. Extrapolating, based on the running time benchmark (see Figure 2A), our GPU-tRNA-mapper requires 160 minutes for this task. On the other hand, parasail requires for the same task over 4000 minutes or close to 3 days, illustrating the benefits of our mapper and allowing for optimization of mapping parameters and testing other random models for the alignment score.

**Fig. 2:**
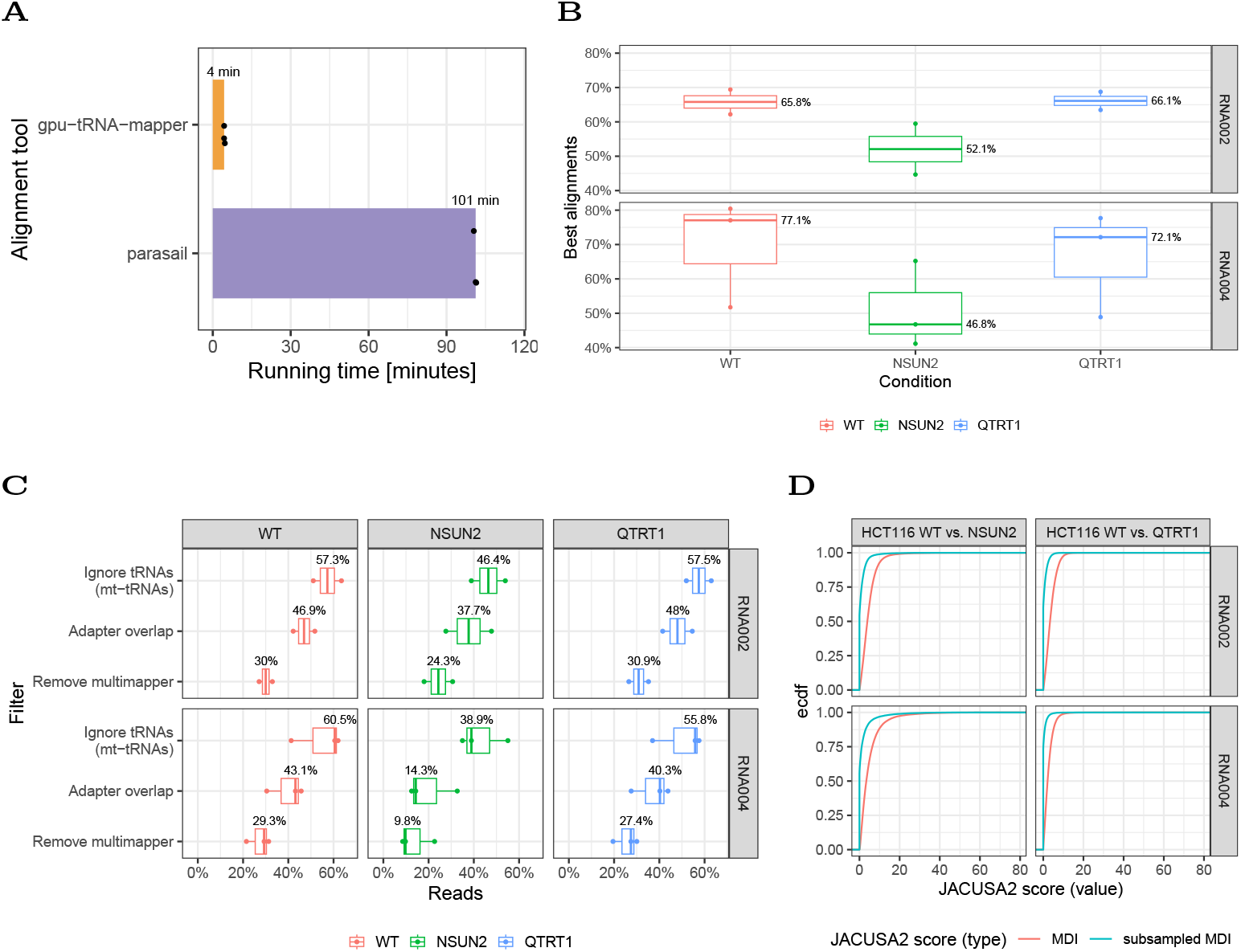
(A) Runtime comparison using a subset of data (10^6^ reads). (B) Percentage of best alignments after mapping, removing random alignments (with the specificity of 99.9%) for HCT116 data sets for chemistry RNA002 and RNA004. (C) Alignment statistics of reads for the HCT116 cell line after each filtering step. (D) Comparison of JACUSA2 score subsampling for HCT116 cell line data sequenced with different chemistries.

#### 2.1.2 Automatic alignment score threshold determination

We estimate the score cutoff for a given specificity level by comparing the distribution of true vs random alignment scores (Figure 1C and Supplementary Figure S1A). If we set the alignment precision to 99.9%, we retain best alignments for between 50 - 70% of all reads in the input (Figure 2B)

#### 2.1.3 Post-alignment filtering steps

We enable some flexibility in downstream processing by offering a set of userconfigurable filters (see Figure 1D): Ignore certain tRNAs from the reference, require a minimal adapter overlap on both ends, and remove multi-mapping reads. Figure 2C summarizes the effect of this series of filtering steps. For example, in the wildtype RNA004 sample, we start with a median read pool size of 77.1 % of library input. Approximately 17% of mt-tRNA reads are removed with the first filter. We required an overlap of 20% for both adapters, which removes another 17% of reads. If we omit reads mapping to multiple reference sequences with the same best score, we decrease the pool size to 29.3 % of the original pool size. Similar effects can be observed for the tRNA RNA002 library (see Figure 2B,C).

#### 2.1.4 Enhancing robustness through JACUSA2 score subsampling

We use JACUSA2 [11] to calculate differences in mismatches, deletions, and insertions (MDI score) between 2 conditions. However, given the higher variability of the data, high MDI scores might already emerge from minor differences when read coverages vary substantially between conditions. To account for this observation, we introduced and implemented an efficient subsampling strategy (see Figure 1E and Supplementary Figure S2). Briefly, the MDI score is normalized by subsampling base calls from the condition with the highest coverage according to the coverage in the other condition (see Supplementary Figure S2 for details). This normalization is performed at the site level, reducing overall scores as shown in Figure 2D.

### 2.2 Analysis of tRNA modifications in the HCT116 cell line

We use human HCT116 cells to create gene knock outs of known tRNA-modifying proteins. Specifically, we generated knock outs for two enzymes, QTRT1 and NSUN2. Both wildtype and knock out cell lines were sequenced on Nanopore sequencing chemistry RNA002 (see Supplementary Figures for 3-7 for results) and RNA004 chemistry, which is the latest iteration of direct RNA sequencing.

The *QTRT1* gene encodes the catalytic subunit of queuine tRNA-ribosyltransferase (TGT), an enzyme complex responsible for the modification of tRNAs. Known targets for QTRT1 mediated tRNA modifications are isoacceptors that carry one of the following amino acids: asparagine, aspartic acid, histidine and tyrosine [13].

The *NSUN2* gene encodes a methyltransferase that catalyzes the methylation of cytosine to 5-methylcytosine (m5C). Furthermore, it is known to stabilize tRNA and prevent its degradation [14].

#### 2.2.1 Analysis of nuclear-tRNAs in QTRT1 and NSUN2 HCT116 knock out cell lines

We sequenced RNA from wildtype, QTRT1 and NSUN2 knock out cell lines with RNA004 chemistry. Results for RNA002 may be found in the Supplementary Figures S3-7. Pairwise QutRNA2 analyses were executed on wildtype vs. QTRT1 samples and wildtype vs. NSUN2 samples (see Methods 5.1). False alignments were removed with a specificity cutoff of 99.9% in a first step after mapping, keeping the best alignments as shown in Figure 2B. Sample specific alignment score distributions and thresholds are shown in Supplementary Figure S1A.

We retained the highest percentage of reads with a median of 77.1% in the wildtype sample, followed by QTRT1 with 72.1%. This number was considerably lower for NSUN2. Here, we had retained best alignments for only 46.8% of reads. The numbers for sequencing with chemistry RNA002 were similar, with NSUN2 having the lowest percentage (52.1%) of best alignments among the three conditions. Wildtype and QTRT1 are on par with ≈ 66% each. When comparing the chemistries, RNA004 shows a higher percentage for wildtype and QTRT1 than for the chemistry RNA002, except for NSUN2. The lower rate of best alignment for NSUN2 might indicate the impact of the enzyme on tRNA stability. The absence of NSUN2 may cause tRNA fragmentation. This is also documented by a shift of read lengths for NSUN2 in comparison to both other conditions (see Supplementary Figure S1B). Subsequently, we filtered all best alignments post hoc to meet certain formal criteria. Figure 2C shows the impact on read counts for this stepwise procedure. First, we remove mt-tRNAs with the “Ignore tRNAs”-filter to focus on nuclear encoded tRNAs. Approximately 16% of sequenced reads in wildtype and QTRT1 are mt-tRNAs and are presented elsewhere downstream. The second filter, “Adapter overlap”, retains reads that have at least 20% overlap with the 5’- and 3’-adapter. This ensures that all remaining reads cover the entire tRNA locus full length. While for wildtype and QTRT1 17% and 15% of reads are filtered, over 24% of NSUN2 reads do not sufficiently cover the adapters. The last filter step removes reads that map to multiple tRNAs with the same best score. Taken together, 29% of the reads remain for the wildtype samples and 27% of the reads for QTRT1 samples. NSUN2 shows the lowest median percentage of remaining reads with ≈ 10% of reads fulfilling the aforementioned requirements. This holds true for both chemistries.

Candidate positions for tRNA modification changes are then identified by calculating subsampled MDI scores (see Figure 2D) for wildtype vs. QTRT1 and wildtype vs. NSUN2 comparisons

As a last step, we translate the reference coordinates from our database to widely accepted Sprinzl coordinates using a eukaryotic-specific covariance model. Normalized JACUSA2 MDI scores for each comparison are presented as heatmaps in Sprinzl coordinates. Known target positions are highlighted by red boxes. Figure 3A depicts the scores calculated for expected substrates of QTRT1, which are tRNAs with GU(N) anticodons, such as tRNA-Asp, -Asn, -His, and -Tyr [15]. The strongest signal was observed for tRNA-Asp-GTC around Sprinzl positions 30 - 35, which overlaps with the expected position 34. Figure 3B shows the same scenario for NSUN2. NSUN2 predominantly targets the cytosine (C) positions 48, 49, and 50 within the variable loop (V-loop) of tRNAs for methylation, a process crucial for maintaining cellular function [16] (see Supplementary Figure S9 for details). Noteably, we observe the highest scoring sites for known positions 48-50 or direct neighbors, such as position 51. NSUN2 adds another m5C modification at position 34 of tRNA-Leu-CAA.

**Fig. 3:**
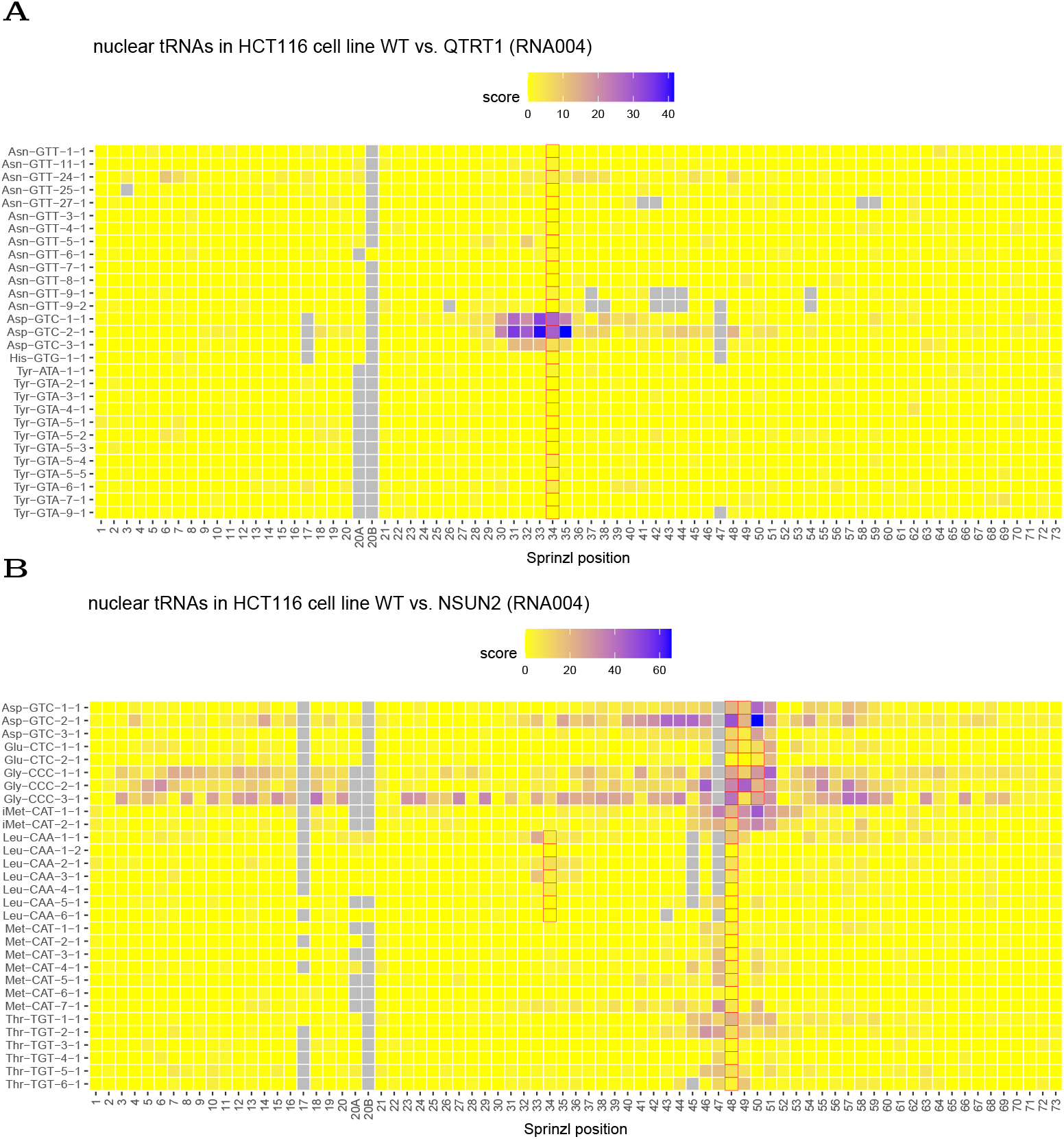
Shown are results for the comparison of HCT116 wildtype to the respective QTRT1 or NSUN2 knock out cell line and subsequently sequenced with RNA004 chemistry. Red boxes indicate the expected modified positions. (A) Wildtype vs. QTRT1. Depicted are subsampled JACUSA2 MDI scores for known tRNA targets: Asn, Asp, His, and Tyr that were covered by at least 30 reads. (B) Scores are shown for wildtype vs. NSUN2 and known targets: Gln, Glu, His, Leu, and Thr (see Supplementary Figure S9 for details.

#### 2.2.2 Analysis of mt-tRNAs in QTRT1 and NSUN2 HCT116 knock out cell lines

We employed an mt-tRNA-specific workflow similar to the one presented in the previous section. This time, we omitted nuclear encoded tRNAs and retained mt-tRNAs instead. The adapter overlap filter leads to a loss of 8-10% of sequenced reads across all conditions. This accounts for almost 50% of the mt-tRNAs (see Supplementary Figure S8B). This loss turned out to be lower for the RNA002 chemistry, where only 25% of mt-tRNAs are removed upon adapter overlap (see Supplementary Figure S6B). We calculated normalized MDI scores for all mt-tRNAs with JACUSA2 and displayed them in heatmaps. Figure 4A shows the scores for the wildtype vs. QTRT1 comparison. We observe two tRNAs with an elevated score at position 34. MT-TH shows the highest score and encodes for aspartate. MT-TY encodes for tyrosine and shows an appreciable yet lower score level at position 34. In Figure 4B, the scores for the wildtype vs. NSUN2 comparison are shown for all mt-tRNAs. Here, the situation is more complex: We observed signals for positions 48, 49 and around positions 43, 44 for some mt-tRNAs. In Van Haute *et al*. [17], it has been shown that for the modification of positions 48, 49, and 50 in mt-tRNAs, NSUN2 is required. When comparing the results to the previous chemistry, similar results are observed for wildtype vs. QTRT1, where MT-TH at position has the strongest signal (see Supplementary Figure S7A). For NSUN2, only modifications in MT-TH at positions 42, 43, 44, 45, and 46 are observed in both chemistries (see Supplementary Figure S7B).

**Fig. 4:**
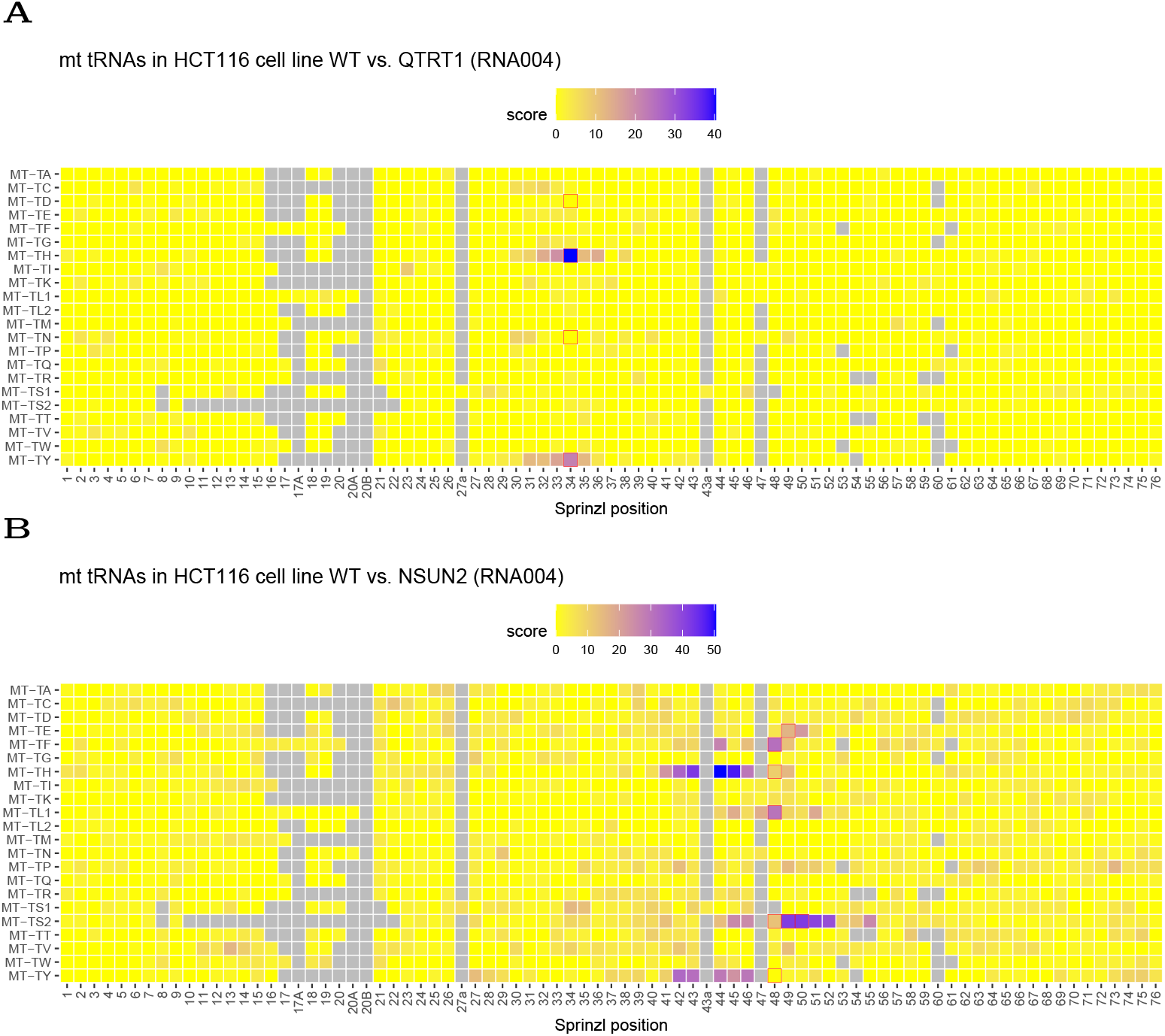
Depicted are subsampled JACUSA2 MDI scores for all mt-tRNAs in HCT116 cells. Red boxes indicate the expected modified positions according to [1]. (A) Wildtype vs. QTRT1 (B) Wildtype vs. NSUN2.

### 2.3 Analysis of nuclear-tRNAs in murine Qtrt1 knock out samples

Finally, we investigated the effects of *Qtrt1* knock out in mice, to exemplify that QutRNA2 can be used to analyze the modification effects in diverse organisms. We use mouse embryonic stem cells (mESC) and mouse hippocampus (mHC) tissue. At the time of sample preparation, RNA002 was the only available chemistry. We used the nuclear-tRNA workflow presented for the HCT116 cell line and analyzed wildtype vs. Qtrt1 knock out in mouse samples mentioned above. See Supplementary Figure S10A for the threshold calculation to filter random alignments. Approximately 71% of sequenced reads could be mapped, passing the random alignment filter when using the mESC (see Supplementary Figures S10B,C for the impact of filters on read length and read counts). As previously outlined, we have identified a set of known Qtrt1 targets.

Figure 5A shows the scores for mESC, and high scores can be observed at the known position 34 for tRNAs carrying asparagine, aspartate, and histidine. Traces of scores at Sprinzl positions 28-35 for tyrosine might indicate tRNA modification. We carried out the same analysis for mHC (see Supplementary Figures S11A,B,C for details on filtering) and achieved similar score profiles (see Figure 5B). Wildtype vs. Qtrt1 in mESC and mHC share the modified tRNAs. Even the tyrosine carrying tRNAs show slightly elevated scores at positions 30-32, much narrower when compared with mESC.

**Fig. 5:**
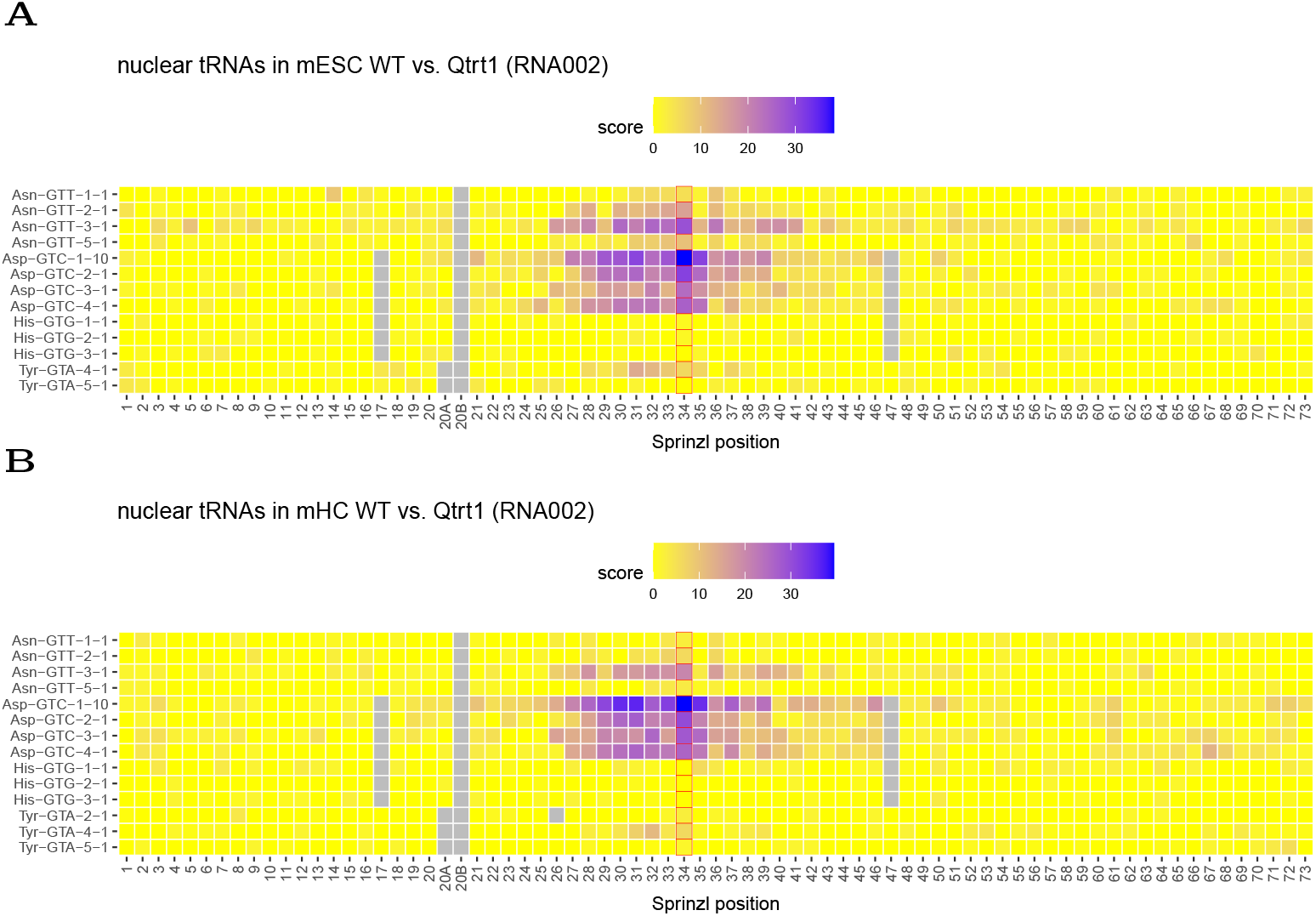
Subsampled JACUSA2 MDI scores wildtype and Qtrt1-deficient mouse ESC (mESC) cells or hippocampus (mHC). Only known tRNA targets are shown: Asn, Asp, His, and Tyr. Samples were sequenced with RNA002 chemistry. (A) Results for deficiency of Qtrt1 in mESC. (B) Results for deficiency of Qtrt1 in mHC.

## 3 Discussion

We have described and presented QutRNA2 as new software for direct tRNA sequencing on the Nanopore platform. To the best of our knowledge, QutRNA2 is unprecedented in its design: It introduces a new rapid GPU-accelerated optimal alignment approach with moderate hardware requirements (e.g. Laptop / Desktop with GPU gaming card suffices). It offers a statistically principled way to reduce the number of false alignments. QutRNA2 introduces a new and more robust scoring scheme to identify candidate positions in complex tRNomes and presents the respective results as heatmaps in widely accepted Sprinzl coordinates.

### 3.1 tRNA modifications in human and mouse samples

We have selected two well known tRNA modifiers, QTRT1 and NSUN2, for a proof-of-principle experiment. The TGT enzyme complex, composed of QTRT1 and QTRT2, specifically recognizes nuclear-encoded tRNAs with GUN anticodons: tRNA-Asp (anticodon GUC), tRNA-Asn (anticodon GUU), tRNA-His (anticodon GUG), tRNA-Tyr (anticodon GUA). Figure 3A shows all tRNA candidates, which fall into this class and meet minimal coverage requirements. We observe a dominant signal for tRNA-Asp (anticodon GUC), but no modification in any other position. We need to keep in mind that this pattern could originate for many reasons: For example, [18] showed that modification rates vary substantially between tRNAs in human cell culture settings with tRNA-Asp being the fastest. This is where we see the strongest signal. Moreover, tRNA-Asp and tRNA-Tyr undergo further glycosylation to mannosyl-queuosine (manQ) and galactosyl-queuosine (galQ), respectively [19]. We need to take this context into account when interpreting our data. However, it is beyond the scope of this software article to investigate this further.

Overall, the data looks noisier for cytoplasmic targets of NSUN2. However, our score profiles are in very good agreement with Figure 1D from Blanco et al. [16] and we capture the previously described m5C modification sites very well.

For Mitochondrial tRNAs, suspected targets of QTRT1 include mt-tRNAs for Asp, Asn, His, or Tyr. Intriguingly, we see a clear modification signal for mt-tRNA-His (MT-TH gene) and mt-tRNA-Tyr (MT-TY gene) in Figure 4A. This would predict different Q34 modifications between cytoplasmic and mitochondrial tRNAs.

### 3.2 Comparison with other methods

A comparison with other methods turned out to be difficult. At the time of writing, none of the peer-reviewed published open source software is capable of running on our data set without heavily modifying the underlying original software. Lucas *et al*. [5] have released a workflow on GitHub that works exclusively on yeast. A more advanced and modern version of this workflow seems to be commercially available from Immagina Biotech (https://www.immaginabiotech.com). Given these circumstances, we did not pursue this option any further.

In another recent publication, Shaw *et al*. [20] present a software for identifying tRNA modifications in yeast using RNA002 chemistry. However, the method requires an IVT sample to train an HMM-based error model for mismatch, insertion and deletion calling, and second, after some inspection, the internal coordinate system seems to be defined only for yeast, making it impossible to compare human or mouse data.

## 4 Conclusion

QutRNA2 is unique in its capabilities. First, we contribute an optimal local alignment approach with 25-fold speedup over our previous approach. This performance increase is useful in optimizing and exploring workflow parameters without sacrificing alignment accuracy. Second, alignments are selected based on their statistical significance and their compliance with formal requirements (e.g., full tRNA coverage). Third, the JACUSA2 component, which identifies candidate positions of differential RNA modification, was enhanced with a subsampling module. This normalization step increases the signal-to-noise ratio and reduces the number of reported candidate sites without sacrificing the detection performance of known modification sites. Moreover, QutRNA2 offers customizable plots to monitor and control the impact of filtering on read length and counts. Finally, we provide an automated mapping of reference sequence coordinates to Sprinzl positions if a matching covariance model is available for the studied organism. In our utility evaluation, we have applied QutRNA2 to data from different organisms, cells, and sequencing chemistries. We have demonstrated that QutRNA2 identifies known targets of QTRT1 and NSUN2 in human HCT116 cells regardless of their genomic origin (nuclear or mitochondrial genome). We also show that analysis results are stable across different sequencing chemistries.

QutRNA2 performed equally well on a different complex tRNA transcriptome from mouse samples, which were either obtained from cell culture or brain tissue.

In summary, QutRNA2 is the only publicly available open source solution that works on high-volume and complex tRNome data sets at the time of writing.

## 5 Methods

### 5.1 HCT116 cell culture

Human HCT116 cell lines were acquired from ATCC and authenticated by Multiplex Human Cell Line Authentication Test (Multiplexion). HCT116 cells were cultured in McCoy’s 5A Medium (Gibco), supplemented with 10% FBS (Gibco), 1% L-glutamine (Gibco), and 1% penicillin/streptomycin (Gibco). HCT116 cells were transiently transfected with the lentiCRISPRv1-sgRNA plasmid (Addgene, USA), targeting the QTRT1 and NSUN2 genes, using Lipofectamine 3000 reagent (Invitrogen, Thermo Fisher Scientific, USA) according to the manufacturer’s instructions. The sgRNAs used were as follows: sgQTRT1 (CACCGGAGCTGATCCAGAAAGCAA), resulting in a homozygous insertion of a C in the QTRT1 gene; and sgNSUN2 (AGCGGCCGGAGGACGCGGAGGATGGCGCCGAGGGTGGT), leading to a homozygous deletion of a G in the NSUN2 gene. After five days of puromycin selection, single clones were isolated and homozygous mutations were confirmed by Sanger sequencing.

### 5.2 Mus musculus cell culture

E14 mESCs were cultured in KnockOut DMEM medium (Gibco). Supplemented with 10% FBS (Gibco), 1% GlutaMAX (Gibco), 1% penicillin/streptomycin (Gibco), 100 μM beta-mercaptoethanol (Sigma-Aldrich), and 1.2 x 103 U LIF (ESGRO), these cells were grown on 0.2% gelatin-coated plates. As previously described in [21], a knock out mESC clone was generated using CRISPR/Cas9 genome editing where the homozygous mutation of a 19 bp deletion in the Qtrt1 gene was confirmed.

#### 5.2.1 Animal handling

The husbandry of mice was performed at the Mannheim Faculty of Medicine, University of Heidelberg, according to applicable laws and regulations. Mice were kept at 12:12 light:dark cycles with standard housing temperature of 18-23°C and tissue dissection was performed in strict compliance with national and international guidelines for the Care and Use of Laboratory Animals (Regierungspräsidium Karlsruhe, Germany). Hippocampal regions from wild type mice and Qtrt1 mutant (Q1; C57BL/6J-Qtrt1em1Tuo) mice [22] were collected.

### 5.3 Total RNA isolation and small RNA purification

Using TRIzol (Invitrogen), total RNA was isolated from either 10^5^-10^7^ cells or approximately 50 mg mouse tissue, according to the manufacturer’s instructions. Prior to adding chloroform, mouse hippocampal tissue went through an additional homogenization, using Polytron PT1200E Homogenizer (Kinematica). The concentration and purity of the total RNA were determined by Nanodrop measurement. Small and large RNA fractions were purified from total RNA using the RNA Clean & Concentrator-25 kit (Zymo Research).

### 5.4 Nanopore direct tRNA sequencing

Nanopore tRNA-sequencing was performed as described by [5] with modifications as described below. First, tRNAs were deacylated by incubation of 2 μg small RNA fraction in 100 mM Tris pH 9.0 for 30 min at 37°C. Deacylated tRNA were then purified using RNA Clean & Concentrator-5 kit (Zymo Research). The tRNA splint adapters were annealed as described [5] with a final concentration of 24 μM and stored at -80°C until usage.

#### 5.4.1 RNA002 library preparation and sequencing

The deacylated tRNA in 10 μl nuclease-free H_2_O was used completely for ligation of the splint adapters. The reaction mixture (50 μl total volume) furthermore contained 2 μl annealed splint adapter, 1x T4 RNA ligase 2 buffer (New England Biolabs), 20% PEG8000 (New England Biolabs), 40 U RiboLock RNase Inhibitor (Thermo Fisher Scientific) and 10 U T4 Rnl2 (New England Biolabs). Reactions were incubated 2 h at room temperature. For clean up, 100 μl RNA Clean XP beads were added (15 min, room temperature, gentle mixing). Beads were washed twice with 200 μl 80% ethanol and resuspended in 10 μl nuclease-free H_2_O. After a 10 min incubation at room temperature, the beads were collected on a magnet and the supernatant transferred to a new tube.

Reverse transcription adapter (RTA), as described by Oxford Nanopore Technologies, was prepared at a final concentration of 8.4 μM. 1 μl of this tRNA-RTA was ligated to the splinted tRNA using T4 DNA ligase. Reactions were assembled in a total volume of 20 μl in 1x Quick ligation buffer with 40 μl RiboLock and 3 kU T4 DNA Ligase (New England Biolabs) and incubated 30 min at room temperature. In parallel, a reverse transcription mixture composed of 8 μl 5x first strand synthesis buffer (Thermo Fisher Scientific), 4 μl 0.1 M DTT, 2 μl 10 mM dNTP (Thermo Fisher Scientific) and 4 μl nuclease-free H_2_O was assembled. After ligation, the reverse transcription mixture was added by gentle pipetting. Reaction was completed by the addition of 2 μl Superscript IV (Thermo Fisher Scientific) and incubated 50 min at 50°C, followed by enzyme denaturation (10 min, 70°C). 80 μl RNA Clean XP beads were added to the reaction and incubated 15 min at room temperature with gentle agitation. Beads were washed twice with 80% ethanol and RNA eluted in 20 μl nuclease-free H_2_O.

6 μl RMX adapter (Oxford Nanopore Technologies, SQK-RNA002) was ligated in a total volume of 40 μl in 1x Quick ligation buffer with 6 kU T4 DNA ligase for 30 min at room temperature. Reactions were cleaned up with 80 μl RNA Clean XP beads (10 min, room temperature) and washed twice with 150 μl WSB (SQK-RNA002). Libraries were eluted in 21 μl ELB (SQK-RNA002, 10 min, room temperature) and the final concentration determined using the Qubit dsDNA HS kit (Thermo Fisher Scientific).

Libraries were loaded completely on MinION R9.4.1 flow cells. Sequencing was started with the bulk file option enabled and 48 h run time. Runs were stopped when no substantial data were generated anymore. The bulk output files were used for sequencing simulations with custom MinKNOW settings as described [5]. Custom runs were performed on MinKNOW 23.07.12 equipped with Dorado 7.1.4 and high accuracy basecalling.

#### 5.4.2 RNA004 library preparation and sequencing

Splint adapter ligation was performed as described for the RNA002 sequencing, however 40 U RNasin (Promega) was used instead of RiboLock. For clean up, 90 μl Magnetic beads for tRNA purification (BioDynami) were used.

RTA ligation was carried out as described for RNA002 sequencing, again with RNasin instead of RiboLock. Reverse transcription was performed with Induro reverse transcriptase (New England Biolabs). A reverse transcription mixture composed of 8 μl 5x Induro buffer, 2 μl 10 mM dNTPs and 8 μl nuclease-free H_2_O was mixed by pipetting with the ligation reaction. Reaction was completed by the addition of 2 μl Induro reverse transcriptase and incubated 30 min at 60°C, followed by enzyme denaturation (10 min, 70°C). Reactions were cleaned up by the addition of 76 μl BioDynami tRNA beads and eluted in 23 μl nuclease-free H_2_O.

The RLA adapter (Oxford Nanopore Technologies, SQK-RNA004) was ligated as described for the RTA adapter (RNA002). Reactions were cleaned up using 72 μl BioDynami tRNA beads and eluted in 33 μl REB (SQK-RNA004) and the final concentration was determined using the Qubit dsDNA HS kit.

Libraries were loaded completely on Promethion RP4 flow cells and sequenced on a P24 A100 device with high-accuracy basecalling (Q score threshold 7) and 5 Gb estimated bases as data target. Due to the higher output and decreased lower read limit (https://community.nanoporetech.com/posts/software-release-24-06-for) run simulations were not used.

### 5.5 tRNA mapping and alignment

The mapping of raw reads from sequencing tRNA libraries was performed by the GPU-accelerated alignment library gpu-tRNA-mapper. First, the best alignment coordinates for each read are called with the GPU. In the final step, the alignments for the precomputed coordinates were extended with parasail v2.6.2 [23], and standard SAM output [24] was produced. gpu-tRNA-mapper was used with the following parameters: *–scoring 1,-1,-1,-1 –batchsize 100000*. All analyses were carried out on a Slurm cluster with diverse nodes. The GPU-accelerated alignment was run on a NVIDIA Quadro RTX 6000.

#### 5.5.1 GPU-accelerated mapping

GPUs are able to execute thousands of threads simultaneously. Our GPU-accelerated alignment functions use the CUDA programming language and follow the general ideas used by CUDASW++4.0 [25]. An alignment between two sequences is processed by a group of up to 32 threads, which can exchange data efficiently via so-called *warp intrinsics*. In contrast to CUDASW++4.0 which provides a GPU-accelerated local alignment with affine gap penalties for proteins, our gpu-tRNA-mapper targets the alphabet A,C,G,T/U,N and provides functions for both local alignments and semiglobal alignments using affine gap penalties. In addition to a score-only computation, we also provide new GPU kernels which find the optimal score as well as the start position and end position of the corresponding traceback path. All alignment functions operate on 32-bit values to avoid inexact alignment scores caused by the limited resolution of narrower data types.

As the first step in the mapping process, the optimal local alignment scores between reads and references are computed on the GPU. Next, we inspect the scores to identify the best matching reference sequences for each read. For those references, we recompute the alignment using the variant which tracks the start and end positions. This concludes the GPU-accelerated portion of the workflow. On the CPU, the start and end positions are then used to compute the optimal alignment traceback using parasail. Last, we write the results to a file in SAM format.

Our software supports NVIDIA GPU architectures with compute capability ≥ 7.0 and VRAM capacity of at least 4GB. It has been tested on Volta, Turing, Ampere, Ada, Hopper, and Blackwell GPUs.

#### 5.5.2 Running time analysis

The running time requirements of parasail and the gpu-tRNA-mapper were studied on a downsampled data set. We used the RNA004 sequencing data set of the HCT116 cell line and downsampled wildtype and QTRT1 samples to 10^6^ reads each. The identical alignment scoring parameters were used for parasail and the gpu-tRNA-mapper. However, modifying batch size and the number of threads did not influence the running time of parasail. In contrast to the gpu-tRNA-mapper, parasail does not calculate the best alignment. We implemented a custom Python script to obtain the highest-scoring alignments for Parasail results. The measured GPU runtimes also include necessary data transfers between GPU and CPU over the PCIe bus.

#### 5.5.3 Human (GRCh38/hg38) reference

The genomic and mature reference sequence information for Homo sapiens (GRCh38 / hg38) tRNAs was downloaded from https://gtrnadb.ucsc.edu [26]. The suffix “withintron” was added to the name of tRNAs containing an intron in the genomic FASTA file. Mature tRNA sequences were converted to DNA and merged with the genomic FASTA file. Finally, the resulting file was filtered by removing tRNAs with identical sequences and retaining the first occurrence of a duplicated sequence. 5’-splint adapter (CCTAAGAGCAAGAAGAAGCCTGGN + tRNA) and 3’-splint adapter (GGCTTCTTCTTGCTCTTAGGAAAAAAAAAA) were added to the reference sequence and used to map raw reads.

#### 5.5.4 Mus musculus (GRCm38/mm10)

Likewise, the sequencing information for Mus musculus (GRCm38/mm10) was obtained from [26], and the procedure as outlined for human was carried out.

### 5.6 Filtering reads

We implemented filters that can be applied in series to remove spurious alignments.

#### 5.6.1 Filter random alignments

This filter will create a random model for alignments by mapping reads to the reversed reference sequence. Then, based on this random alignment model and a precision defined by the user, an alignment threshold is calculated and applied to filter reads. The precision was set to 99.9%. Furthermore, an informative plot is created that presents a summary of the distributions of sequenced samples and alignment scores.

### 5.7 Adapter overlap

This filter retains reads overlapping with the 5’ and 3’-splint adapters. We required an overlap of 20% for both adapters.

### 5.8 Remove multimappers

We used this filter to remove multimapping reads that align more than once with the same alignment score at different tRNAs.

### 5.9 JACUSA2 scoring of mismatches and INDELs

We used an extended version of JACUSA2 [11] to score and distinguish modified and unmodified bases regarding reference and non-reference base calls or INDEL calls at a candidate site. The MDI score consists of a linear combination of test statistics that measure the difference in mismatches (M), deletions (D), and insertions (I) when comparing two experimental conditions [12]. The modified version (v2.1.15) of JACUSA2 has been called with the following parameters: *-f X:reads:nonref ratio:insertion ratio:deletion ratio -u DirMult:subsampleRuns=100 -m 1 -q 1 -D -i -P1 FR-SECONDSTRAND -P2 FR-SECONDSTRAND*.

#### 5.9.1 Subsampling of scores

In case of uneven coverage across libraries and conditions, we have increased the signal-to-noise ratio in the JACUSA2 statistics by a subsampling approach (see Supplementary Figure S2 for details). In brief, raw MDI scores are normalized by subtracting the maximum of subsampled scores. Subsampled scores are calculated by subsampling reads from the condition with the highest coverage, according to the coverage of the other condition. We used 100 subsamplings (-u DirMult:subsampleRuns=100) to calculate subsampled MDI scores for each site.

### 5.10 Visualization of tRNA modifications

Calculated JACUSA2 scores were visualized in heatmaps where rows represent the tRNAs and columns depict the exact Sprinzl coordinate. Where necessary, plots of subsampled MDI scores were restricted to show specific tRNAs or mark positions of special interest. All plots were generated with the parameters *-ignore trnas=withintron, – remove prefix = tRNA-, –hide varm*. The number of minimal reads per replicate was set to *–min reads=30* or *–min reads=50*.

#### 5.10.1 Secondary structure alignment

Covariance models for eukaryotic tRNA were obtained from https://github.com/UCSC-LoweLab/tRAX/blob/master/TRNAinf-euk.cm [27]. tRNAs of each species were split into nuclear and mitochondrial tRNAs, and the matching covariance model along with Infernal 1.1.5 [28] was used to calculate secondary structure predictions for nuclear-tRNAs with the following parameters: *–notrunc– nonbanded -g*. The sequence to Sprinzl coordinate mapping was obtained from [1]. The naming scheme from [10] labeled the consensus secondary structure.

## Supplementary information

We provide Supplementary Text, Figures and Tables in PDF format.

## Declarations

### Ethics approval and consent to participate

Not applicable

### Consent for publication

Not applicable

### Availability of data and materials

The score code for QutRNA2 will be made available at: https://github.com/dieterichlab/QutRNA2. A Conda based installation procedure is described there too. The score code for the gpu-tRNA-mapper will be made available at: https://github.com/fkallen/gpu-tRNA-mapper.

### Authors’ contributions

MP implemented the entire workflow, added enhancements to existing components, analyzed the data, generated figures and wrote the manuscript. WG performed cell culture and animal experiments and provided total RNA extracts and wrote parts of the methods section. INV performed all Nanopore sequencing experiments, analyzed data and wrote the manuscript. FK and BS implemented the GPU-accelerated mapper. FT designed and supervised the experimental part. BS supervised the study. CD conceived and designed the approach, analyzed data, helped with debugging, supervised the study and wrote the paper. All authors read and approved the final manuscript.

## Acknowledgements

We are grateful to Harald Wilhelmi for his excellent assistance in supporting software development and running and maintaining the Dieterich Lab compute cluster.

