## Supplementary Materials for "QutRNA2: Robust tRNA modification discovery from Nanopore direct tRNA sequencing"

### Supplementary Material for QutRNA2: Robust tRNA modification discovery from Nanopore direct tRNA sequencing

October 20, 2025

### Supplementary Figures

#### Results for HCT116 nuclear-tRNAs (RNA004)

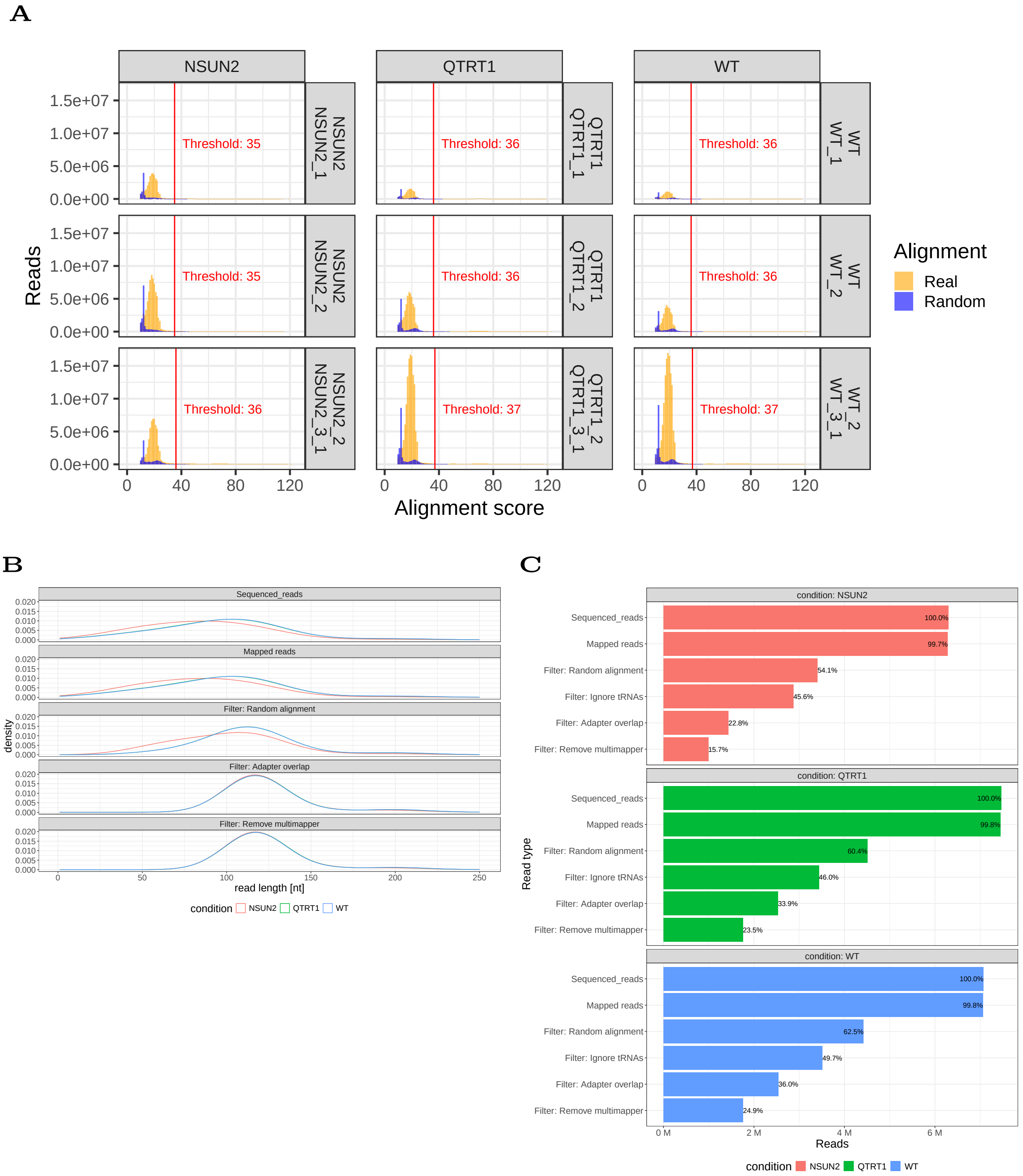


Figure 1: QutRNA2 intermediate results for HCT116 (RNA004) nuclear-tRNAs. (A) Calculated alignment score thresholds for each subsample with precision set to 99.9%. (B) Impact of subsequent filtering steps on read length for each condition. (C) same for Read counts.

##### New subsampling algorithm in JACUSA2

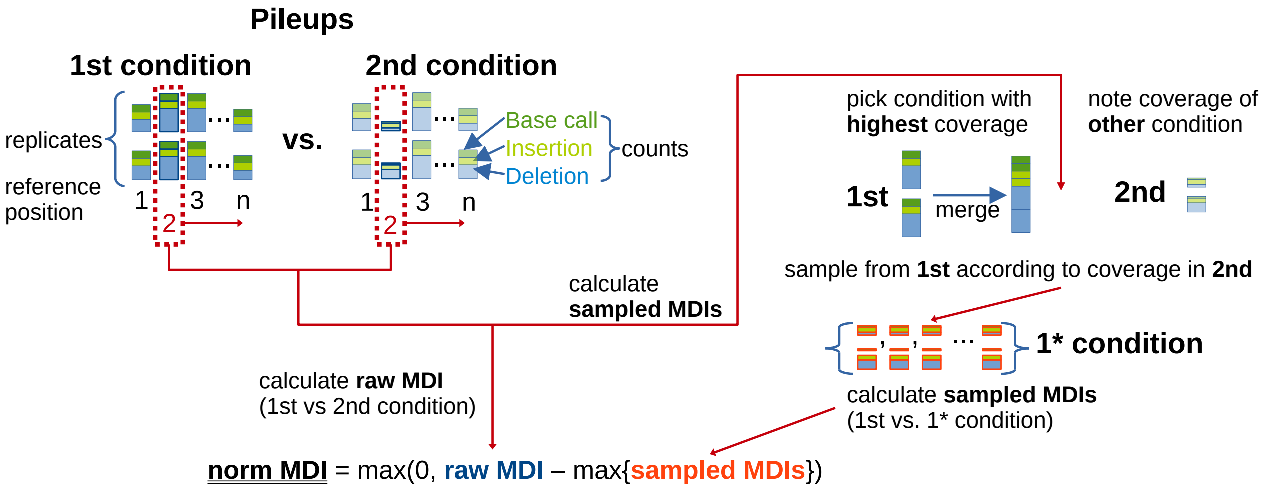


Figure 2: Illustration of subsampling from read stacks/pileups in JACUSA2 to improve signal-to-noise ratio at positions with different read coverages. Raw JACUSA2 scores are calculated from pileups of both conditions. Raw scores are normalized by subtracting the maximum of subsampled scores. Subsampling ca be repeated multiple times, and the user can set the number of runs. Subsampled scores are calculated by subsampling reads from the condition with the highest coverage, according to the coverage of the other condition.

#### Filtering results for HCT116 nuclear tRNAs (RNA002)

**A**

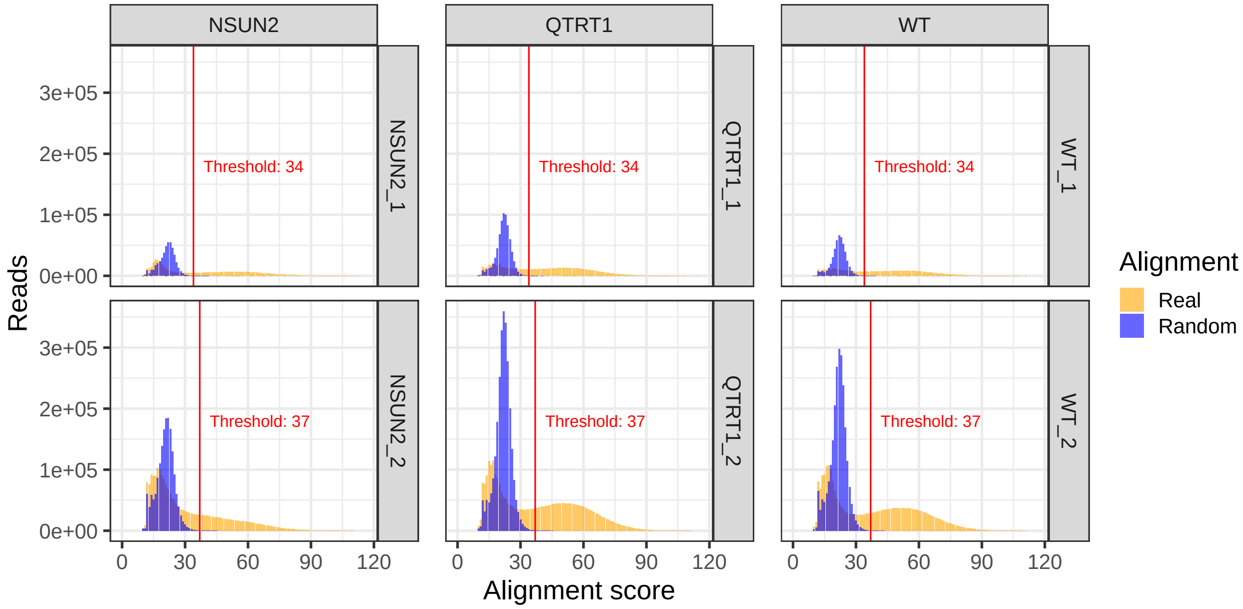


**B**

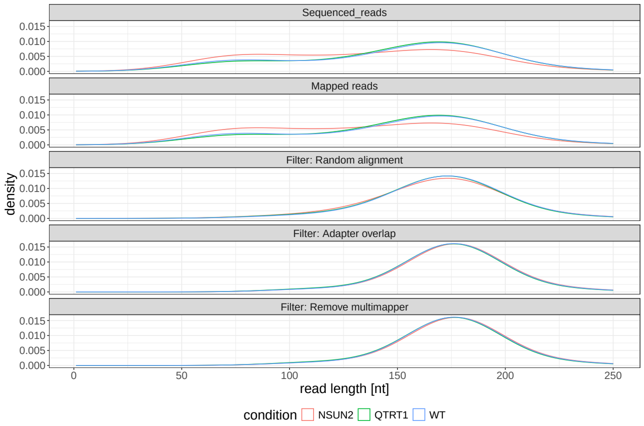


Figure 3: Results for HCT116 (RNA002) nuclear-tRNAs. (A) Calculated alignment score thresholds for each subsample. (B) Filtering effects on read length for each condition.

**A**

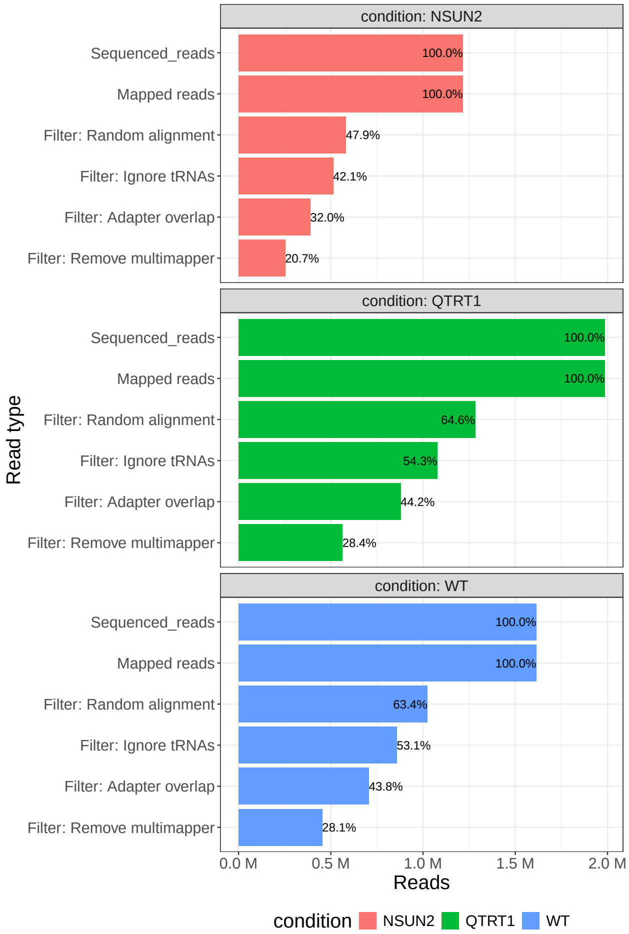


Figure 4: Results for HCT116 (RNA002) nuclear-tRNAs. Filtering effects on Read counts.

### Extended results for HCT116 nuclear-tRNAs (RNA002)

**A**

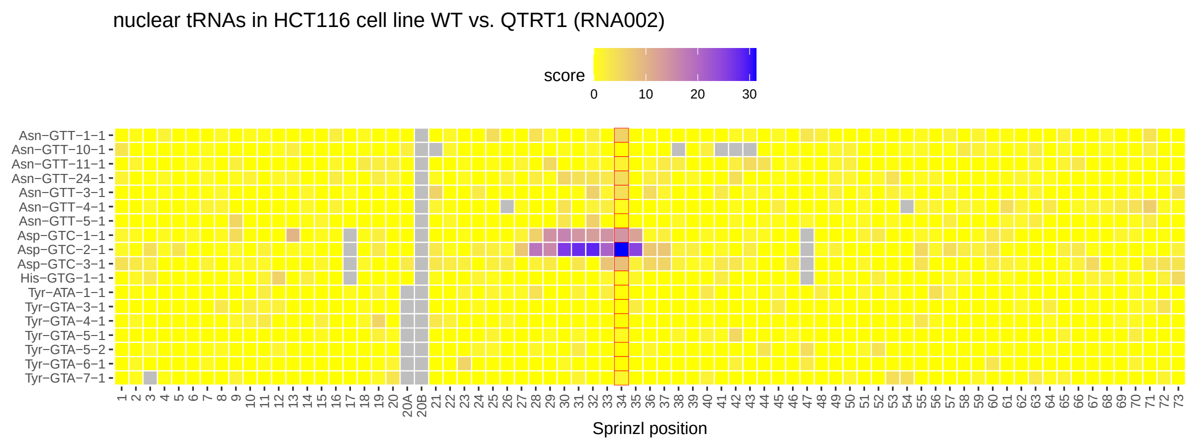


**B**

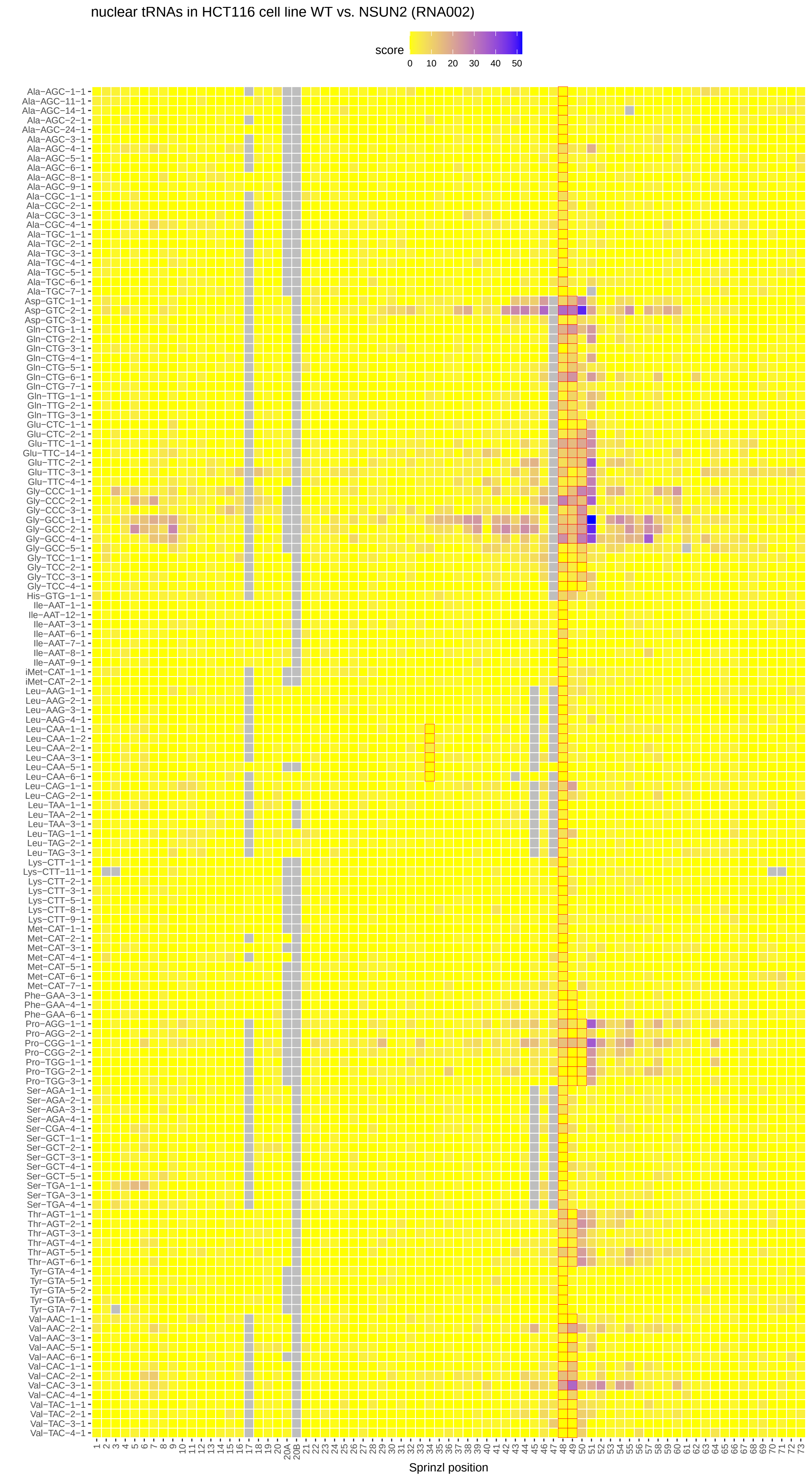


Figure 5: Comparison of wildtype vs. NSUN2 and QTRT1 knock outs in HCT116 cells (RNA002 chemistry). (A) Subsampled MDI Scores for condition wildtype vs. QTRT1 in relevant tRNAs (Asn, Asp, His, and Tyr) are shown with a coverage of at least 30 reads. (B) Scores are shown for wildtype vs. NSUN2. Complete list of potential NSUN2 targets as defined in Figure 1B in [1]. Only tRNAs with at least 50 reads in each replicate are shown.

#### Filtering results for HCT116 mt-tRNAs (RNA002)

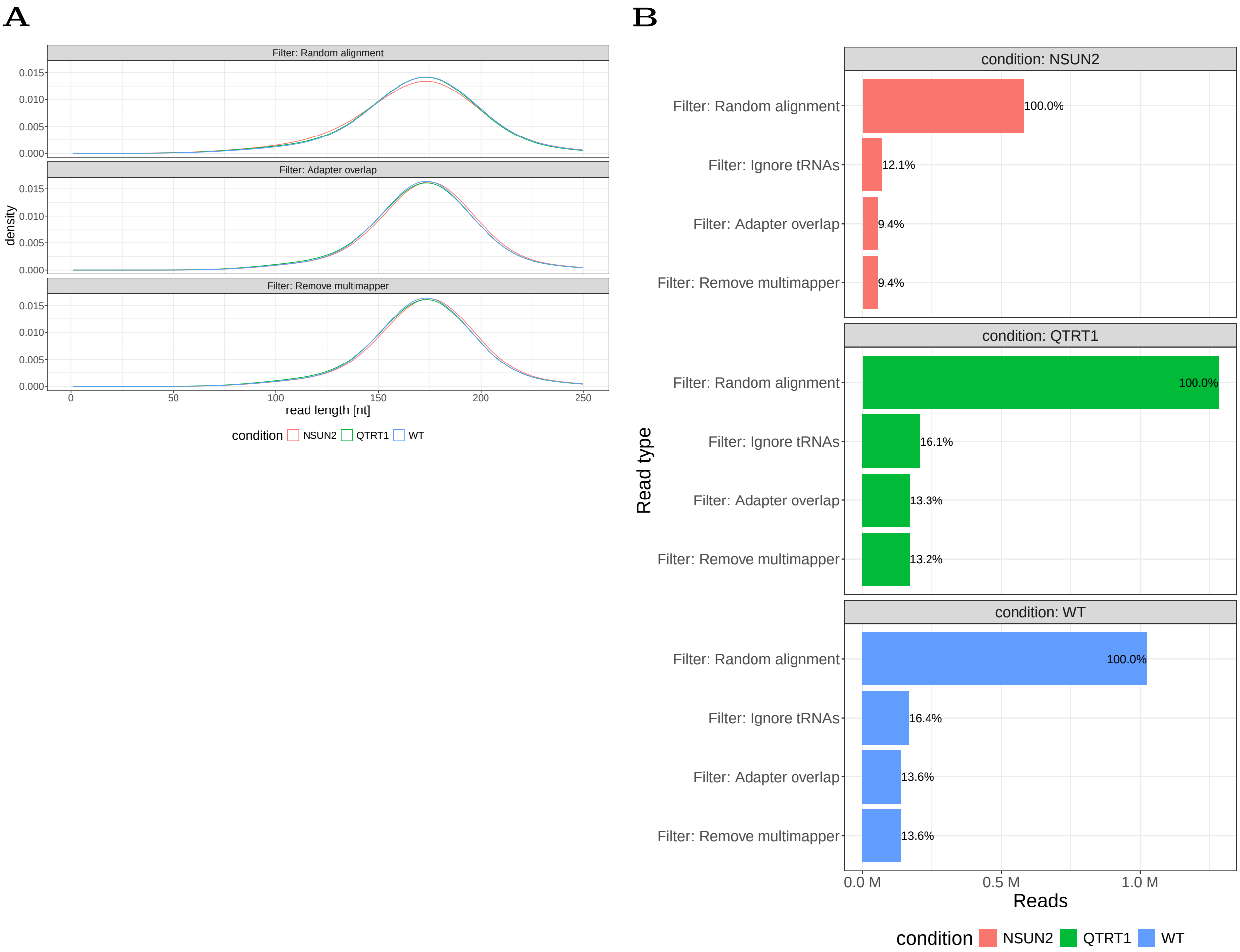


Figure 6: Detailed results for mt-tRNAs for the HCT116 cell line (RNA002). Reads were taken from the "Ignore tRNAs" filtering step of the initial mapping. (A) Distribution of read lengths for mt-tRNAs. (B) Distribution of read counts for mt-tRNAs.

### Results for HCT116 mt-tRNAs (RNA002)

## A

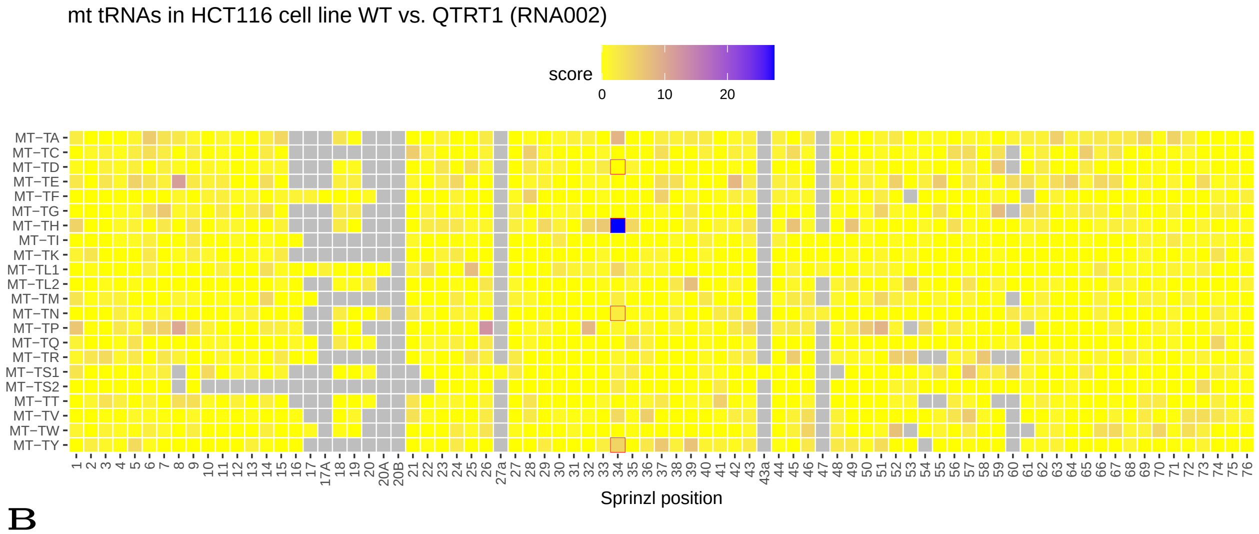


## B

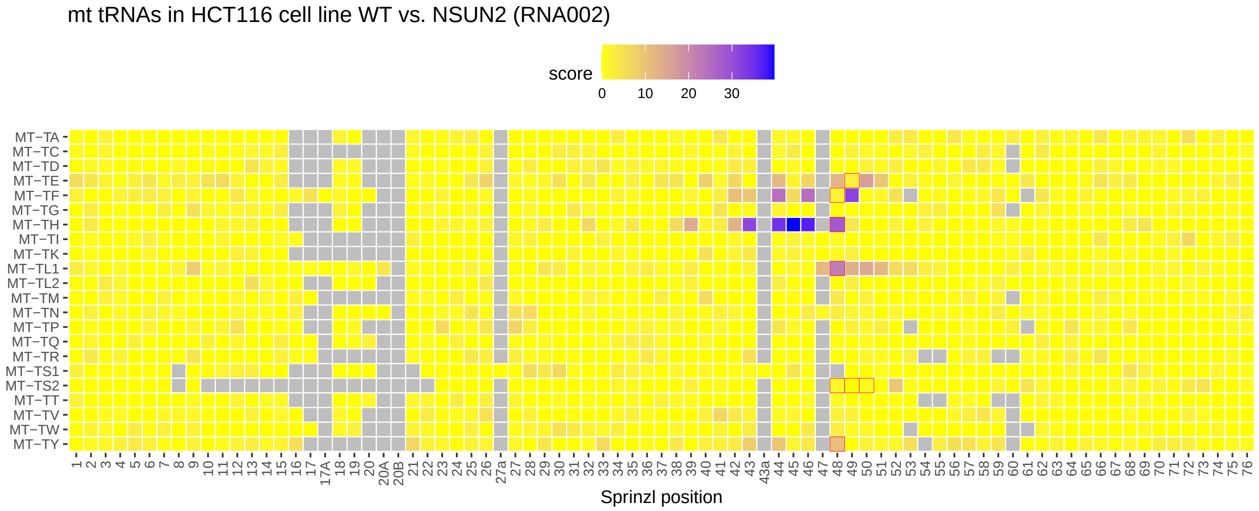


Figure 7: Comparison of wildtype vs. NSUN2 and QTRT1 knock outs in HCT116 cells sequenced with RNA002 chemistry. Displayed are subsampled MDI scores for all mt-tRNAs. Red boxes indicate tRNAs for the following amino acids: Asn, Asp, His, and Tyr. (A) Scores for wildtype vs. QTRT1. (B) Scores for wildtype vs. NSUN2.

### Filtering results for HCT116 mt-tRNAs (RNA004)

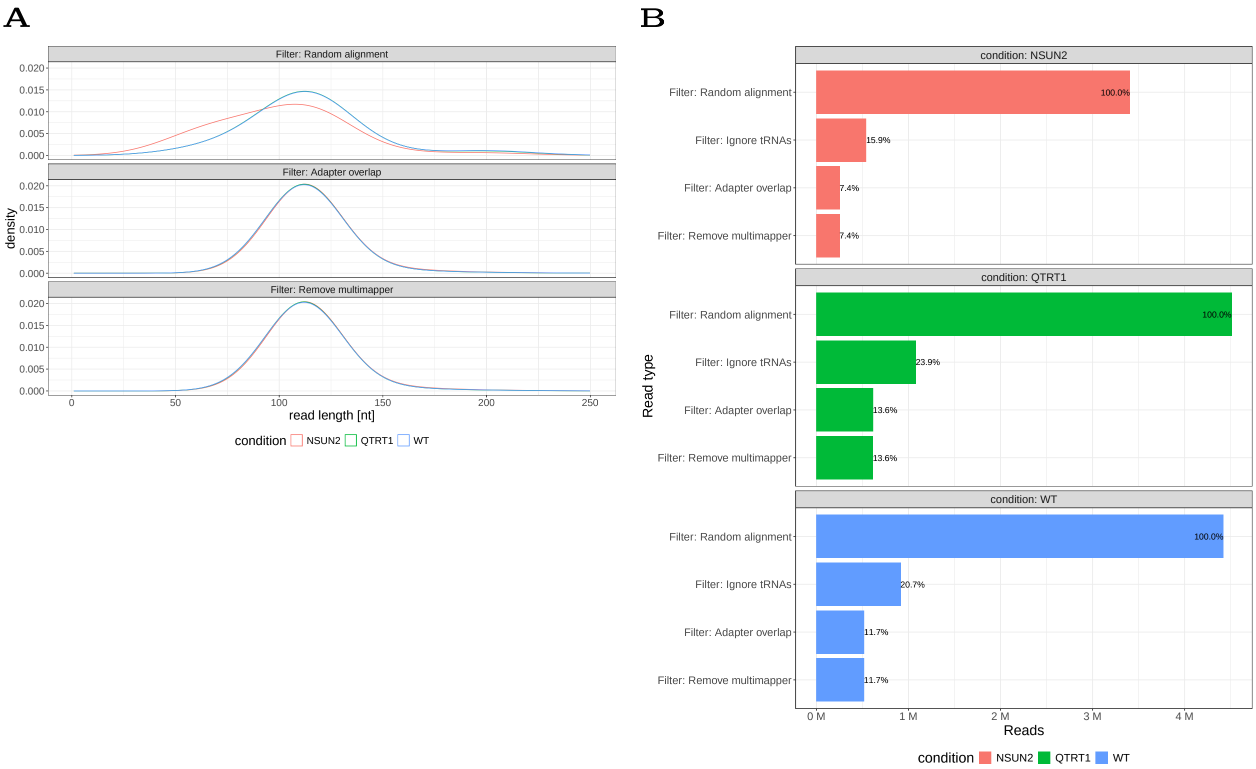


Figure 8: Detailed results for mt-tRNAs for the HCT116 cell line (RNA004). Reads were taken from the "Ignore tRNAs" filtering step of the initial mapping. (A) Distribution of read lengths for mt-tRNAs. (B) Distribution of read counts for mt-tRNAs.

#### Extended results for HCT116 nuclear-tRNAs (RNA004)

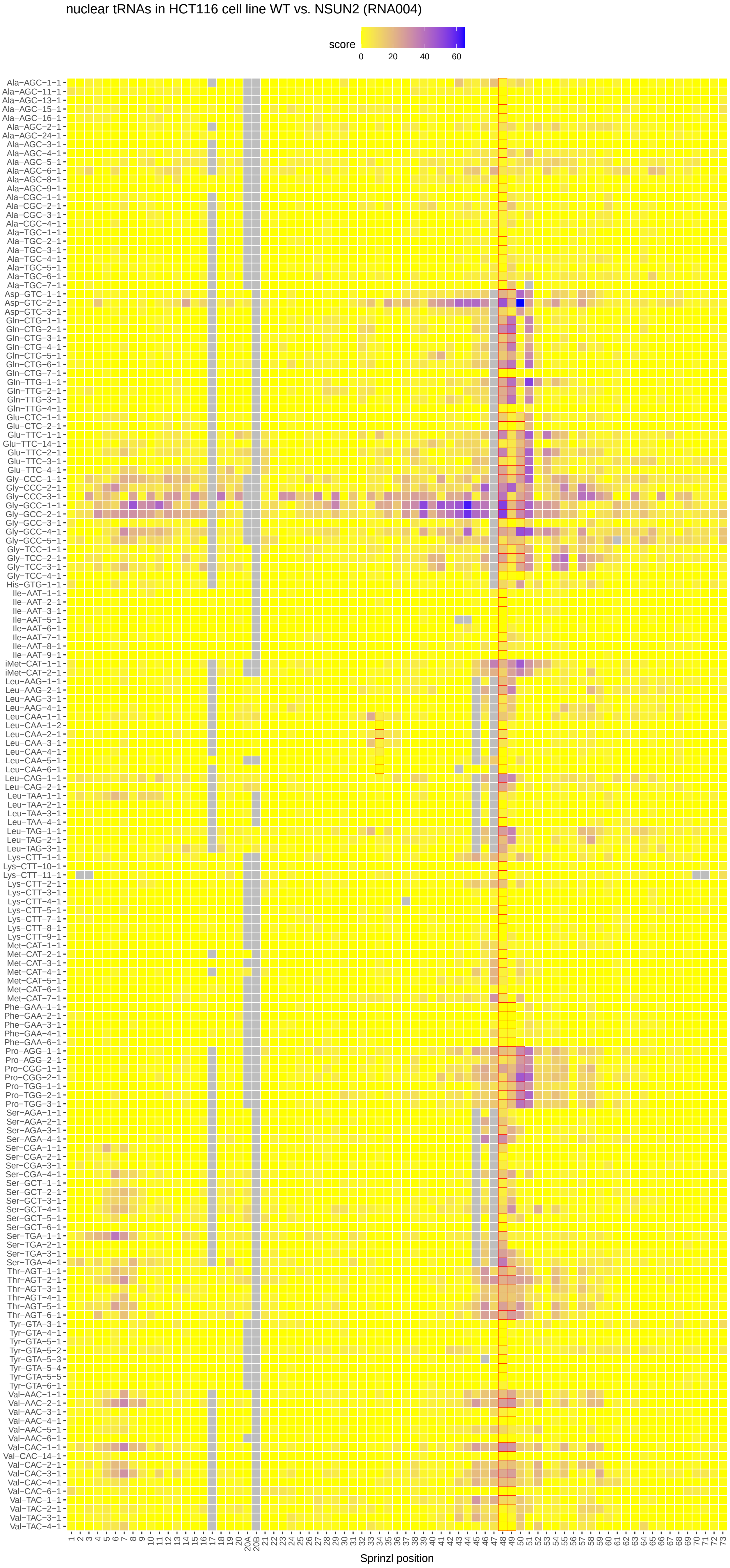


Figure 9: Results for HCT116 nuclear-tRNAs (RNA004) for wildtype vs. NSUN2. Complete list of potential NSUN2 targets as defined in Figure 1B in [1]. Only tRNAs with 50 ≥ reads in each replicate are shown.

#### Filtering results for mESC nuclear-tRNAs (RNA002)

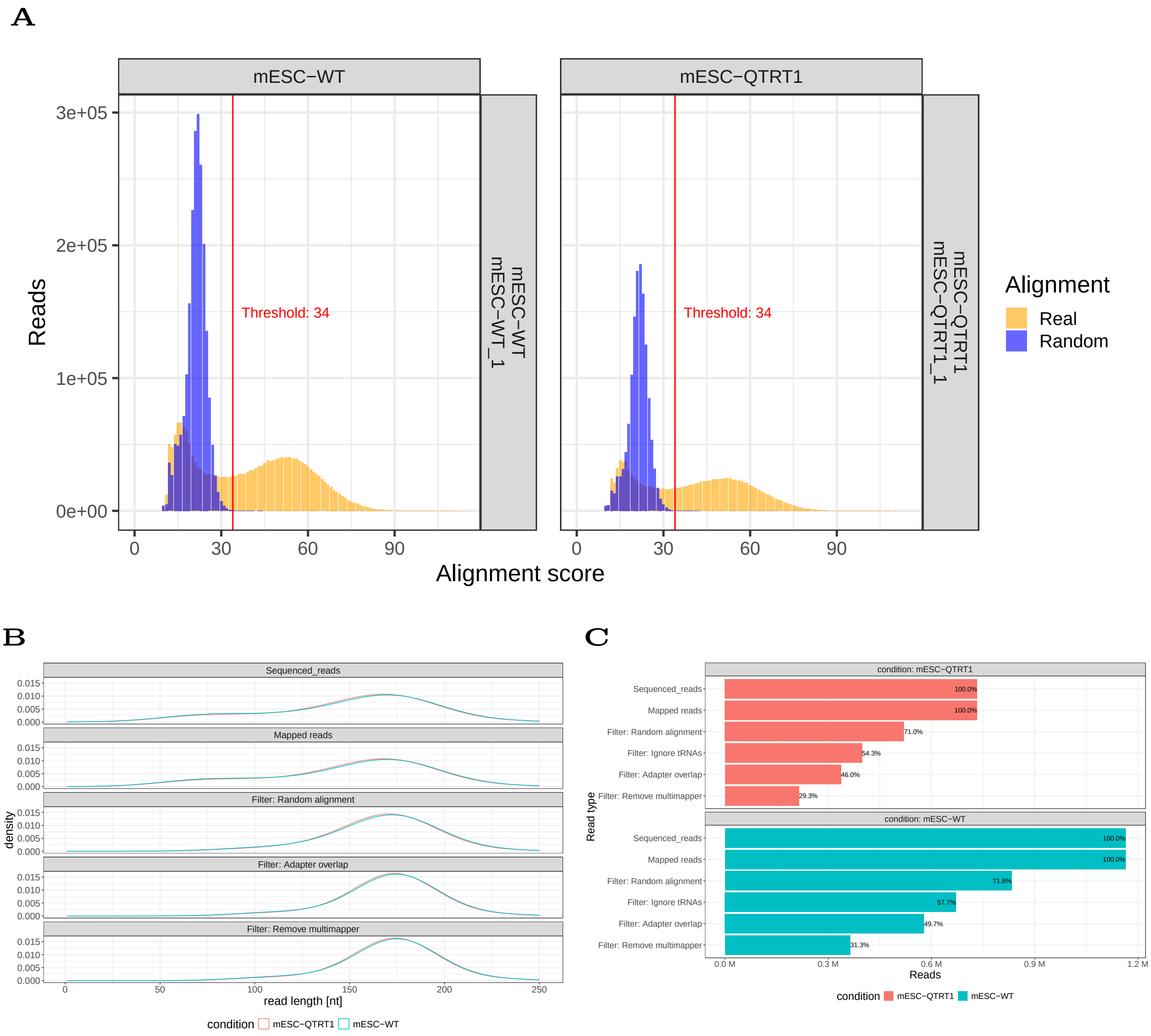


Figure 10: Results for mESC (RNA002) nuclear-tRNAs. (A) Calculated alignment score thresholds for each sample. (B) Filtering effects on read length for each condition. (C) Filtering effects on Read counts.

#### Filtering results for mHC nuclear-tRNAs (RNA002)

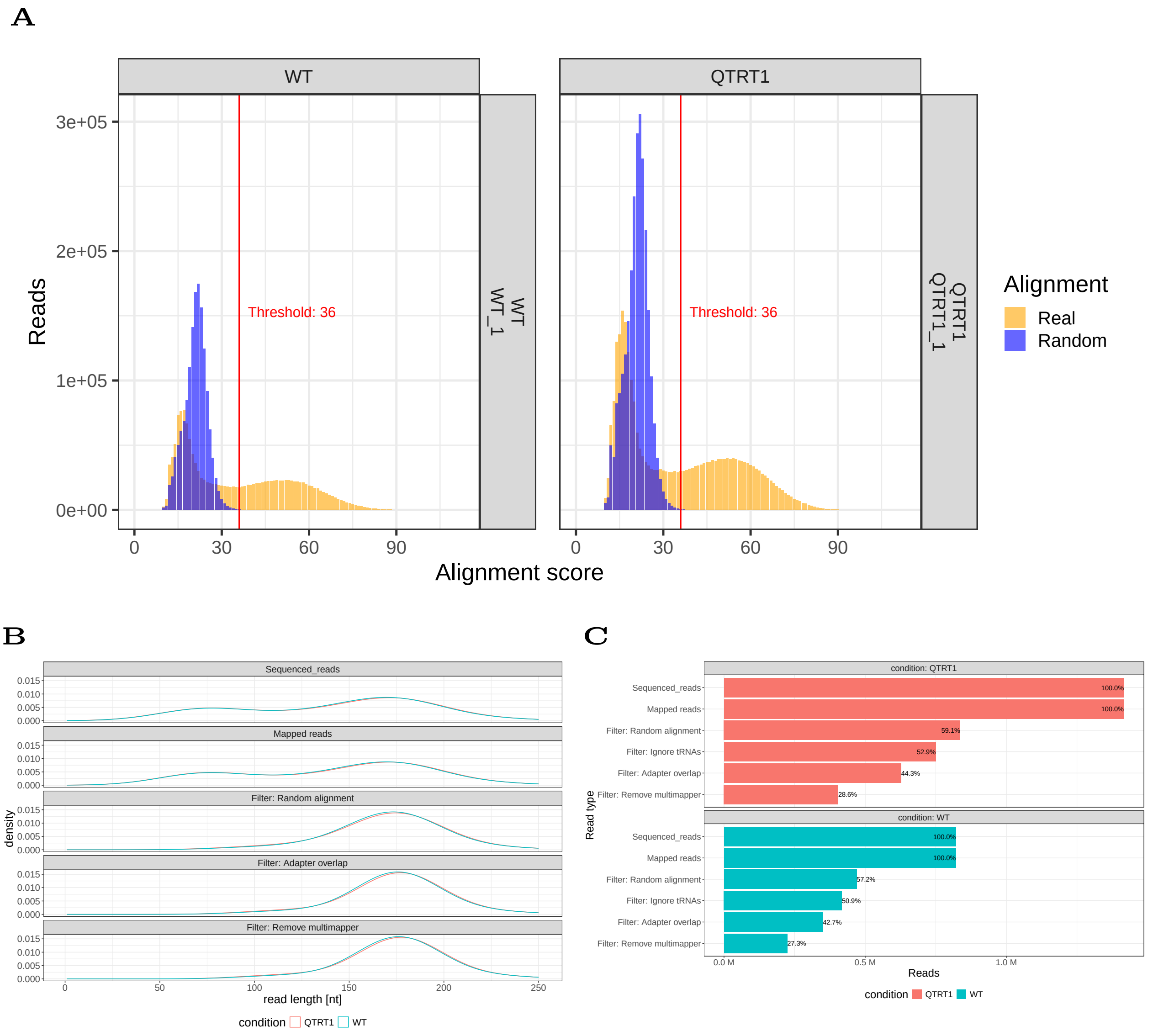


Figure 11: Results for mHC (RNA002) nuclear-tRNAs. (A) Calculated alignment score thresholds for each sample. (B) Filtering effects on read length for each condition. (C) Filtering effects on Read counts.

### Supplementary Tables

Description of supplementary tables below.

#### Supplementary Table 1: Sequencing data generated for this study

| Identifier | cell line/ tissue | genotype | replicate | chemistry |
|---|---|---|---|---|
| 662eb444 | HCT116 | wildtype | biological 1 | RNA002 |
| 0a96bd04 | HCT116 | wildtype | biological 2 | RNA002 |
| 2ebcd7ff | HCT116 | QTRT1 KO | biological 1 | RNA002 |
| b9a81c30 | HCT116 | QTRT1 KO | biological 2 | RNA002 |
| fba868e1 | HCT116 | NSUN2 KO | biological 1 | RNA002 |
| 57c0bbfe | HCT116 | NSUN2 KO | biological 2 | RNA002 |
| e377bc82 | HCT116 | wildtype | biological 1, technical 1 | RNA004 |
| 79f237fe | HCT116 | wildtype | biological 1, technical 2 | RNA004 |
| 92e99007 | HCT116 | wildtype | biological 2 | RNA004 |
| 074dfc6d | HCT116 | QTRT1 KO | biological 1, technical 1 | RNA004 |
| a9cd5076 | HCT116 | QTRT1 KO | biological 1, technical 2 | RNA004 |
| da529cd9 | HCT116 | QTRT1 KO | biological 2 | RNA004 |
| 2b29a8ab | HCT116 | NSUN2 KO | biological 1, technical 1 | RNA004 |
| 3d823b92 | HCT116 | NSUN2 KO | biological 1, technical 2 | RNA004 |
| 06149b48 | HCT116 | NSUN2 KO | biological 2 | RNA004 |
| cc9edd48 | mESC | wildtype | - | RNA002 |
| 6e750df8 | mESC | Qtrt1 KO | - | RNA002 |
| 1d8e08a0 | mouse hippocampus | wildtype | - | RNA002 |
| dbd5437e | mouse hippocampus | Qtrt1 KO | - | RNA002 |

Table 1: Sequencing data generated for this study
